# Cross-Species Adaptation of RETFound for Rodent OCT Age Estimation Reveals Strong CNN Baselines in Data-Scarce Space Biology

**DOI:** 10.64898/2026.04.22.720210

**Authors:** Alireza Hayati, Jian Gong, Vaishnavi Nagesh, Pinar Avci, Ariel Yuhan Ong, Mouayad Masalkhi, Justin Engelmann, Fathi Karouia, Pearse A. Keane, Sylvain V. Costes, Lauren M. Sanders

**Affiliations:** Students Research Committee (SRC), Qazvin University of Medical Sciences, Qazvin, Iran; Independent Researcher; School of Computing, University of Wyoming, Laramie, WY, USA; School of Medicine, University College Dublin, Belfield, Dublin, Ireland; NIHR Biomedical Research Centre at Moorfields Eye Hospital NHS Foundation Trust, London, UK; Institute of Ophthalmology, University College London, London, UK; Blue Marble Space Institute of Science and Exobiology Branch, NASA Ames Research Center, Moffett Field, CA, USA; Center for Space Biomanufacturing, Synthetic Biology and Digital Health; Trivedi Institute for Space and Global Biomedicine; and Departments of Computational and Systems Biology and Radiation Oncology, University of Pittsburgh School of Medicine, Pittsburgh, PA, USA; Blue Marble Space Institute of Science, Seattle, WA, USA

**Keywords:** Optical Coherence Tomography, Foundation Models, Vision Transformers, Convolutional Neural Networks, Low-Rank Adaptation, Space Biology, Rodent

## Abstract

Space-biology imaging studies are often constrained by severe data scarcity, limiting the development of robust machine-learning biomarkers. Rodent spaceflight and space-analog datasets provide an important preclinical setting for testing transfer-learning strategies, but the extent to which human retinal foundation models can generalize to rodent optical coherence tomography (OCT) remains unclear. Here, we benchmark cross-species adaptation of RETFound, a human retinal Vision Transformer pretrained on 1.6 million retinal images, for chronological age prediction from Brown Norway rat OCT B-scans in the NASA Open Science Data Repository dataset OSD-679.

We adapted RETFound using Low-Rank Adaptation (LoRA) and evaluated performance on control animals under matched 3-fold rat-level cross-validation. We compared RETFound+LoRA with a strong ImageNet-pretrained Xception baseline under matched protocols and included a scratch/random ViT as a supplementary negative-control architecture check. Metrics included mean absolute error (MAE), R^2^, and inter-eye mean absolute difference (MAD).

RETFound+LoRA achieved MAE = 26.20 ± 5.03 days with R^2^ = 0.744 ± 0.049. However, Xception performed better in the primary benchmark (MAE = 19.01 ± 7.67 days, R^2^ = 0.853 ± 0.082), and the matched-fold comparison favored Xception, although this result should be interpreted cautiously given the small number of folds. Inter-eye consistency was maintained across the matched control evaluation, and saliency maps localized model attention to anatomically plausible inner retinal regions.

Together, these results show that human retinal foundation models can transfer to rodent OCT in a scientifically useful way, but also that strong CNN baselines may outperform transformer-based models in small-sample cross-species settings. This preprint provides a reproducible benchmark and baseline framework for future retinal biomarker development in space biology.

**Significance Statement:** Space-biology imaging studies are intrinsically data-limited. This preprint provides a reproducible cross-species benchmark for adapting Earth-trained retinal models to rodent OCT under small-sample conditions, highlights the value of strong CNN baselines, and offers a reusable starting point for future retinal biomarker development in space-relevant datasets.

## 1. Introduction

Aging is a complex, multifactorial phenomenon characterized by the progressive decline of biological systems. Understanding—and ultimately mitigating—this decline requires studying how organisms respond to diverse stressors, including extreme environments such as spaceflight.[1] Space missions and space-analog studies are increasingly utilized as platforms to probe aging-relevant pathways (e.g., genome instability, inflammation, immune dysregulation, and metabolic stress) in both humans and model systems.[2] However, the development of data-driven biomarkers[2] for space biology is constrained by extreme data scarcity. Astronaut cohorts are historically restricted in sample size,[3] highly selected for health, predominantly male,[4] and their data is protected by stringent privacy controls.[5] These factors severely limit the development of space-relevant machine learning models using conventional supervised learning approaches.

In parallel, advances in self-supervised learning have enabled foundation models—large-scale neural networks pretrained on massive, domain-specific datasets—to improve generalization when labeled data are limited. RETFound, a Vision Transformer (ViT) pretrained via masked autoencoding on 1.6 million human retinal images from the UK Biobank and Moorfields Eye Hospital, exemplifies this paradigm.[6] RETFound has shown strong transfer performance across multiple downstream ophthalmic tasks, including diabetic retinopathy and glaucoma detection, prognosis of age-related macular degeneration, and “oculomics” tasks such as prediction of ischemic stroke, myocardial infarction, and Parkinson’s disease.[6] This asymmetry—abundant Earth-based retinal data versus scarce space-based data—motivates a novel strategy: leveraging Earth-trained, human-centric foundation models to enable analysis in data-scarce space biology contexts.

The brown rat (Rattus norvegicus) has traditionally served as a valuable model organism for interspecies knowledge transfer. While lacking a fovea and being rod-dominant, its retina shares the same ten-layered structure as humans,[7,8] and its response to simulated microgravity (via hindlimb suspension, HLS) mirrors key symptoms of Spaceflight-Associated Neuro-Ocular Syndrome (SANS), including optic nerve head remodeling and altered intraocular pressure.[9– 12] Yet, no study has investigated whether a human-trained foundation model like RETFound can be adapted to interpret rat OCT scans for basic phenotypic prediction tasks. Moreover, documenting the steps and benchmarks for such domain adaptation can provide valuable lessons as foundation models are increasingly considered for space biology research. Prior work has applied deep learning to rodent OCT primarily for pathology classification and layer segmentation in experimental models, rather than for cross-species transfer or normative biomarker development, highlighting the gap addressed by the present study.[23,24]

Here, we use data from the NASA GeneLab Open Science Dataset OSD-679, focusing on rodent OCT B-scans collected under normative (control) conditions. We selected chronological age prediction as a benchmark task because it provides a well-defined test of whether retinal structure contains transferable age-related signal across species. In this setting, age prediction serves as a practical first step for evaluating whether a human retinal foundation model can be adapted to rodent OCT under substantial domain shift and limited sample size. Establishing this baseline is important before pursuing more biologically interpretive downstream analyses, such as stress-associated deviations from normative retinal aging, which have been proposed as candidate biomarkers of systemic aging and health outcomes in human retinal imaging studies.[15,16,22] To address the dual challenges of human-to-rodent transfer and data scarcity, we fine-tune RETFound using Low-Rank Adaptation (LoRA), a parameter-efficient strategy that freezes the pretrained model weights and learns task-specific updates through small trainable low-rank matrices inserted into targeted layers. Conceptually, LoRA constrains adaptation to a low-dimensional subspace, reducing the number of trainable parameters while helping preserve the general retinal representations acquired during large-scale self-supervised pretraining.

In this preprint, we present a reproducible cross-species benchmark for retinal age estimation from rodent OCT in a data-scarce space-biology setting. Using OSD-679 control animals, we adapt RETFound with LoRA, compare it against matched CNN and random-transformer baselines, assess inter-eye consistency, and examine saliency localization. Our aim is not only to test whether a human retinal foundation model can transfer to rodent OCT, but also to establish a practical baseline for future stress-associated retinal biomarker studies in space-relevant datasets.

## 2. Methods

### 2.1 Dataset and Experimental Design

This study leveraged the NASA GeneLab Open Science Dataset OSD-679, a ground-based spaceflight-analog experiment that includes longitudinal retinal OCT imaging from Brown Norway rats (Rattus norvegicus). The full dataset comprises retinal OCT images stored across multiple OCT-derived formats. In this work, we focused on the BScanThumb and REGAVG formats, which together represented the majority of usable OCT data. After restricting the dataset to Control animals from Cohorts 1-3 and to the BScanThumb and REGAVG image types, the filtered dataset contained 3,466 B-scans from 123 unique rats (253 rat-eye pairs). Primary modeling focused on Day 0 and Day 90, which together contributed 2,238 B-scans from 123 unique rats (246 rat-eye pairs). Training was performed at the B-scan level, all train/validation/test splits were performed at the rat-ID level, and rat-level predictions were aggregated by late fusion where indicated. The filtered age range therefore spanned 90-360 days (approximately 3-12 months), which corresponds to an early-to-mid-life window within a typical Brown Norway rat lifespan of roughly 2-3 years.

BScanThumb images correspond to single-frame retinal OCT B-scans exported as downsampled thumbnail representations, preserving retinal layer contrast while reducing file size. REGAVG images represent motion-corrected, registered, and averaged B-scans generated by the acquisition system to improve signal-to-noise ratio and structural clarity.

The dataset consists of four experimental cohorts differing in age, sex, and environmental exposure; these cohort definitions were inherited from the original OSD-679 study design and were not created for the present analysis. Among the included cohorts, Cohort 1 corresponded to young males, Cohort 2 to young females, and Cohort 3 to older males. Cohorts 1 and 2 contributed the largest number of images, whereas Cohort 4 contained substantially fewer samples. Due to this pronounced imbalance, Cohort 4 was excluded from downstream modeling. Cohort 3, despite having fewer samples than Cohorts 1 and 2, was retained because it provided a wider chronological age range relevant to age prediction analysis.

For this study, we included only Control animals, defined here as animals housed under standard vivarium conditions without hindlimb suspension in the original OSD-679 experiment. This restriction allowed us to establish a normative baseline for age prediction performance without confounding effects from experimental stressors. A summary of cohort characteristics and sample counts is provided in Supplementary Table S1. Across the primary Day 0 and Day 90 analysis window, Cohort 1 contributed 800 scans from 41 rats, Cohort 2 contributed 1,016 scans from 50 rats, and Cohort 3 contributed 422 scans from 32 rats.

OCT imaging was performed longitudinally at predefined experimental days (Day 0, 7, 14, 28, 90, 97, 114, 128, and 180), spanning a total experimental duration of 180 days. Although the dataset is longitudinal, image and sample counts varied substantially across acquisition days. The majority of images were concentrated at Day 0 and Day 90, whereas several intermediate timepoints contained markedly fewer animals per cohort. Consequently, all primary modeling focused on Day 0 and Day 90, which together accounted for the majority of the filtered dataset (1,478 scans at Day 0 and 760 scans at Day 90) and provided the most stable cohort coverage. These refer to experimental study days rather than days of life.

### 2.2 Image Preprocessing and Augmentation

Raw OCT B-scans were acquired as single-channel (grayscale) images. To ensure compatibility with the pretrained Vision Transformer backbone used in RETFound, each B-scan was converted to a three-channel input by duplicating the grayscale channel. Images were then square-padded to preserve the original aspect ratio, resized to 224 × 224 pixels (the standard RETFound input resolution), and normalized using ImageNet mean and standard deviation.

During training, we applied data augmentation using mild intensity policies. Augmentation included random rotations (±15°), random resized crops (scale 0.9-1.0), horizontal flips (probability 0.5), and mild Gaussian blur (σ 0.0-0.5). Photometric augmentations (color jitter, gamma adjustment) were disabled to preserve OCT-specific intensity relationships. No additional manual image-quality exclusion was applied before model development beyond modality/day/group filtering; instead, quality-related failure modes were examined qualitatively in the worst-case review analyses. Augmentation parameters were selected via Optuna-based hyperparameter optimization (Supplementary Table S8).

### 2.3 Data Splits and Cross-Validation

All data splits were performed at the rat-ID level to prevent any leakage across timepoints or between eyes. We used 3-fold cross-validation (K = 3) on Control animals only to balance robustness with the limited sample size. Using a larger number of folds would have produced very small validation subsets, making the validation loss highly unstable and potentially biasing model selection toward idiosyncratic splits rather than generalizable performance.

For external-holdout backbone ablation experiments, the filtered Day 0/90 dataset contained 248 rats overall (123 Controls and 125 HLS animals). The fixed split used 73 control rats for training, 25 control rats for validation, 25 control rats for held-out control testing, and 125 HLS rats as an external stress/OOD evaluation set, with no rat overlap between sets. Although this HLS external/OOD set was reserved during development and supplementary benchmarking, those results were not a focus of the present control-benchmark paper and are therefore not emphasized in the main narrative. Cross-validation split definitions for the control-only folds are provided in Supplementary Table S9.

For clarity, the training unit in the primary benchmark was the individual OCT B-scan, whereas the split unit was the rat ID. The primary reporting unit was the pooled out-of-fold control eye-day prediction set from the matched 3-fold benchmark. Accordingly, Table 1 summarizes pooled out-of-fold control performance across all eligible eye-day observations, Table 2 reports the same pooled out-of-fold predictions stratified by cohort and study day, and inter-eye MAD was computed as the absolute OD-OS prediction difference for matched rat-day pairs available within the evaluation set.

**Table 1:** RETFound+LoRA Control Performance Under Matched 3-Fold Cross-Validation.

| Metric | Mean ± SD | Fold 0 | Fold 1 | Fold 2 |
| --- | --- | --- | --- | --- |
| MAE (days) | 26.20 ± 5.03 | 31.79 | 22.02 | 24.81 |
| RMSE (days) | 43.24 ± 5.38 | 49.20 | 38.74 | 41.79 |
| R<sup>2</sup> | 0.744 ± 0.049 | 0.688 | 0.775 | 0.770 |
| Pearson r | 0.868 ± 0.024 | 0.840 | 0.881 | 0.882 |

Note: Metrics correspond to pooled out-of-fold control evaluation under the matched 3-fold rat-level cross-validation protocol using the RETFound backbone with LoRA adaptation.

**Table 2:** RETFound+LoRA Control Performance by Cohort and Day (Pooled Out-of-Fold Evaluation)

| Cohort / pooled evaluation | Day 0 MAE | Day 90 MAE | Overall MAE |
| --- | --- | --- | --- |
| Cohort 1 (Young Males; n=41 rats) | 12.51 | 44.10 | 24.83 |
| Cohort 2 (Young Females; n=50 rats) | 4.74 | 22.78 | 11.70 |
| Cohort 3 (Older Males; n=32 rats) | 32.86 | 96.75 | 54.41 |
| Pooled out-of-fold controls | 14.35 | 45.74 | 26.20 |

Note: MAEs in Table 2 are calculated on pooled out-of-fold control eye-day predictions from the matched 3-fold cross-validation protocol. Cohort labels indicate the rat cohorts represented in the Day 0/90 subset; the row values themselves are not late-fusion rat-level averages.

### 2.4 Hyperparameter Optimization

Hyperparameters were selected using Optuna-based optimization on the Control-only training set under 3-fold cross-validation. The search space included optimization settings (e.g., learning rate and weight decay), LoRA configuration, regularization parameters (e.g., dropout), and the categorical choice of augmentation intensity (mild/medium/high). The objective was to minimize the cross-validated validation loss (Smooth L1 / Huber). The final configuration used for all reported experiments corresponds to the best-performing Optuna trial and is summarized in Supplementary Table S8.

### 2.5 Model Architecture

RETFound (a masked-autoencoder Vision Transformer pretrained on 1.6 million human retinal images, including color fundus photographs and OCT B-scans) was adapted for rodent OCT-based age regression. The transformer backbone was frozen throughout training to preserve pretrained retinal representations.

We performed parameter-efficient domain adaptation using Low-Rank Adaptation (LoRA) modules inserted into the self-attention projection layers of the transformer. LoRA hyperparameters (rank = 16, alpha = 32, dropout = 0.1) were selected via Optuna-based optimization. We used LoRA because RETFound is a pretrained retinal foundation model intended for downstream adaptation, whereas the OSD-679 rodent OCT dataset is relatively small and involves substantial domain shift from the human retinal images used during pretraining. LoRA freezes the pretrained backbone and learns task-specific low-rank updates within the transformer, thereby reducing the number of trainable parameters relative to full fine-tuning while preserving access to pretrained retinal representations. This makes LoRA a principled adaptation strategy for a data-scarce space-analog setting, where full end-to-end optimization of a large transformer would be more prone to overfitting and computational inefficiency [3,6,13,20].

A lightweight regression head mapped patch embeddings to a scalar age prediction using global average pooling followed by a 2-layer MLP (512 → 256 → 1 output) with ReLU activations and dropout (p = 0.1).

### 2.6 Training Strategy

The model was trained to regress chronological age (in days) using Control animals only, enabling the learning of a normative retinal aging representation. Optimization was conducted at the per-image level, where individual OCT B-scans served as independent training samples.

To mitigate age imbalance within Controls (younger vs. older animals), we used a weighted random sampler. The optimizer (AdamW, lr = 1×10^−4^, weight decay = 1×10^−4^), loss function (Smooth L1 / Huber), and training schedule (cosine annealing with 5-epoch warmup, 50 epochs total, early stopping patience = 10) were selected through Optuna.

For backbone ablation comparisons, we included Xception (ImageNet-pretrained) as a strong CNN baseline and Random ViT (scratch initialization) as a transformer control. Xception is a well-established convolutional architecture based on depthwise separable convolutions and has precedent in retinal age prediction studies, making it a principled comparator for testing whether retinal foundation-model pretraining provides added value in a small-sample cross-species setting. All ablation models were trained with identical head architectures, hyperparameters, and data splits to ensure fair comparison. [14–16]

### 2.7 Evaluation Metrics

Model performance was assessed using mean absolute error (MAE; primary accuracy metric, in days), root mean square error (RMSE; sensitivity to large errors), coefficient of determination (R^2^), Pearson correlation coefficient (r; reported descriptively and interpreted cautiously in this small-sample setting), and inter-eye mean absolute difference (MAD; internal consistency between OD and OS predictions, in days).

### 2.8 Explainability and Saliency Analysis

Model interpretability was derived directly from the spatial age activation maps produced by the regression head. Activation maps were upsampled to higher spatial resolution (448 × 448 pixels), normalized using percentile clipping (2–98%), smoothed with a Gaussian kernel (σ = 1.0), and overlaid onto the corresponding OCT images with low opacity (α = 0.25). The default visualization colormap was viridis. Saliency outputs were presented as side-by-side panels comprising the original OCT image, the overlay, and the raw activation map.

### 2.9 Statistical Analysis

Cross-validation results are reported as mean ± standard deviation across folds. Cohort-specific and day-specific breakdowns are presented descriptively for transparency. For the prespecified primary backbone comparison, the matched-fold comparison favored Xception (two-sided paired t-test on fold MAE, p = 0.0422), although this inferential result should be interpreted cautiously given the small number of folds. Because only three matched folds were available, fold-based confidence intervals were not emphasized because they would not be stable or especially informative, and the remaining model, cohort, and day comparisons are interpreted descriptively rather than as confirmatory significance tests. Pearson correlation coefficients are likewise reported descriptively. All analyses were conducted in Python 3.9+ using PyTorch 2.0+, scikit-learn, and pandas. Code and configuration files are available at: https://github.com/Alirezahayatimedtech/LoRaRetAgePred

## 3. Results

### 3.1 Model Performance on Control Animals

The LoRA-adapted RETFound model was trained exclusively on 123 Control rats (2,238 B-scans in the Day 0/90 analysis window) to learn normative retinal age prediction. Unless otherwise stated, all primary performance numbers reported in the main text refer to this matched rat-level 3-fold cross-validation benchmark. Under that protocol, RETFound achieved MAE = 26.20 ± 5.03 days, RMSE = 43.24 ± 5.38 days, R^2^ = 0.744 ± 0.049, and Pearson r = 0.868 ± 0.024 on control data (Table 1).

Across folds, the model demonstrated stable performance (Table 1). When pooled across out-of-fold control predictions, the resulting MAE corresponded to 15.8% of the mean evaluated age, indicating that the adapted model captured a substantial fraction of chronological-age variation despite the limited sample size and cross-species transfer setting.

Performance differed across timepoints and cohorts (Table 2). Error was lower at Day 0 than at Day 90, indicating that age prediction was more difficult at older chronological ages in this dataset. We interpret this cautiously: retinal aging is structurally heterogeneous across layers, and age-prediction models often show larger errors near the extremes of the age distribution, both of which may contribute to the observed pattern. [17,18]

Performance also varied by cohort. Cohort 2 (young females) showed the most favorable error profile across both timepoints, whereas Cohort 3 showed the largest error overall and the steepest deterioration at Day 90. In the pooled out-of-fold control evaluation, Cohort 3 entered the study at an older baseline age (270 days), so its Day 0 and Day 90 rows correspond to 270- and 360-day evaluations rather than the 90- and 180-day regime represented by Cohorts 1 and 2. Although sex-associated retinal structural differences have been reported in human OCT studies,[19] we avoid over-interpreting the present pattern as a simple sex effect because this dataset does not disentangle sex from age range, cohort composition, sample-size imbalance, or image-quality variation. Cohort-level differences may therefore reflect multiple overlapping sources of variation rather than a single biological factor.

### 3.2 Backbone Ablation: RETFound vs. Xception vs. Random ViT

To isolate the effect of backbone architecture from pipeline complexity, we conducted a fair ablation study with identical regression heads (GAP + 2-layer MLP), frozen backbones, and matched hyperparameters across three architectures: RETFound (ViT, human-pretrained), Xception (CNN, ImageNet-pretrained), and Random ViT (ViT, scratch initialization).

Results are summarized in Table 3, with the full three-backbone benchmark including the scratch/random transformer reported in Supplementary Table S4. Under matched 3-fold cross-validation, the Xception baseline outperformed RETFound+LoRA on prediction accuracy, showing lower MAE/RMSE and higher R^2^. Because Xception is a strong pretrained CNN with efficient depthwise separable convolutions and prior use in retinal age prediction studies, we view this as an informative benchmark result rather than a comparison against a weak baseline. [14–16]

**Table 3:** Primary Backbone Comparison on Control Data (Matched 3-Fold CV)

| Backbone | MAE (days) | RMSE (days) | R<sup>2</sup> | Inter-Eye MAD (days) |
| --- | --- | --- | --- | --- |
| RETFound + LoRA | 26.20 ± 5.03 | 43.24 ± 5.38 | 0.744 ± 0.049 | 14.22 |
| Xception + GAP | 19.01 ± 7.67 | 32.26 ± 9.89 | 0.853 ± 0.082 | 16.71 |

Xception outperformed RETFound by 7.19 days MAE (27.4% relative improvement) and achieved higher R^2^ (0.853 ± 0.082 vs. 0.744 ± 0.049). The matched-fold comparison favored Xception (paired t-test on fold MAE, p = 0.0422), although this inferential result should be interpreted cautiously given the small number of folds. In the full supplementary benchmark, the scratch/random ViT performed worst overall (MAE = 71.06 ± 5.08 days; R^2^ = 0.027 ± 0.103), confirming the importance of pretrained representations in this cross-species task.

Inter-eye MAD was lower for RETFound (14.22 days) than for Xception (16.71 days). Importantly, the scratch/random ViT showed artificially low inter-eye disagreement in the supplementary benchmark despite very poor age-prediction accuracy, which is more consistent with underfitting or partially collapsed predictions than with anatomically faithful agreement.

### 3.3 LoRA vs. Full Fine-Tuning vs. Frozen Head-Only

We compared three adaptation strategies to justify our parameter-efficient approach: (1) LoRA adapters with frozen backbone, (2) full fine-tuning of all parameters, and (3) frozen backbone with trainable head only.

Results are summarized in Supplementary Table S5. Under the current control-focused protocol, full fine-tuning achieved the best raw control accuracy (control MAE = 26.87 days; R^2^ = 0.662), LoRA produced control MAE = 34.90 days (R^2^ = 0.582), and frozen head-only training degraded markedly to MAE = 55.26 days (R^2^ = 0.264). These adaptation-ablation values come from a separate supplementary protocol and should not be compared directly with the primary matched 3-fold benchmark values in Table 1. LoRA nevertheless remained substantially more parameter-efficient and reached its best validation loss earlier.

LoRA converged faster than full fine-tuning (28 vs. 35 epochs to best validation loss), but it did not achieve the lowest final control MAE in this ablation. The frozen backbone with head-only training performed worst (control MAE = 55.26 days), confirming that some backbone adaptation is necessary for cross-species transfer.

### 3.4 Feature Distillation Ablation (Human vs. Rat-Adapted Teacher)

To evaluate whether feature-level knowledge transfer could improve cross-species adaptation, we performed feature distillation from two teachers to an Xception student: (1) human-pretrained RETFound, and (2) rat-adapted RETFound (MAE-pretrained on 7,073 unlabeled rat OCT images spanning Controls and HLS animals across Cohorts 1-3).

Results are summarized in Supplementary Table S6. To avoid confusion, the Xception value of 23.76 days reported in this paragraph is not the main paper benchmark and should not be compared directly with the primary Xception result of 19.01 ± 7.67 days in Table 3. The 19.01 ± 7.67 value comes from the matched rat-level 3-fold control benchmark used for the main paper, whereas the 23.76 value is the undistilled Xception baseline from a separate supplementary distillation protocol. Within that separate distillation protocol, direct feature distillation from human RETFound (α = 0.3) degraded Xception performance from 23.76 to 26.06 days, suggesting substantial domain mismatch. Using a rat-adapted RETFound teacher (α = 0.1) reduced this penalty to 24.76 days and improved RMSE/R^2^, indicating better tail-error control. However, inter-eye consistency remained worse than the corresponding undistilled Xception baseline under that protocol, and Day 0 accuracy was slightly reduced. These results suggest that while rat-adapted features contain useful signals, direct feature alignment introduces trade-offs that do not justify added complexity for this task.

### 3.5 Inter-Eye Reliability Analysis

Model reliability was assessed using inter-eye mean absolute difference (MAD) between left and right eye predictions. Across the RETFound 3-fold control evaluation, the mean inter-eye difference was 14.22 days, corresponding to 8.56% of mean age.

Inter-eye consistency was preserved across the control evaluation sets. Detailed results are provided in Table 4, supporting internal model stability and anatomical consistency. In the matched 3-fold control comparison, RETFound showed lower inter-eye MAD (14.22 days) than Xception (16.71 days).

**Table 4:** Inter-Eye Reliability in the OSD-679 Control Arm.

| Model / evaluation set | Mean MAD (days) | Median MAD | Q95 MAD | Max MAD | % of Mean Age |
| --- | --- | --- | --- | --- | --- |
| RETFound + LoRA (Control arm, 3-fold out-of-fold) | 14.22 | 7.62 | 48.45 | 81.03 | 8.56% |
| Xception + GAP (Control arm, 3-fold out-of-fold) | 16.71 | 8.02 | 59.69 | 86.19 | 10.07% |

### 3.6 Saliency Map Analysis

Saliency maps (examples in Figure 2) highlighted anatomically meaningful retinal regions and demonstrated consistent spatial localization across folds, supporting non-random model behavior. Dominant activations clustered around the retinal nerve fiber layer (RNFL), optic nerve head-adjacent tissue, and inner retinal layers. This pattern is biologically plausible because these compartments are implicated in age-related retinal remodeling and in spaceflight-relevant neuro-ocular change hypotheses derived from rodent and human analog studies.[9-12]

**Figure 1:**
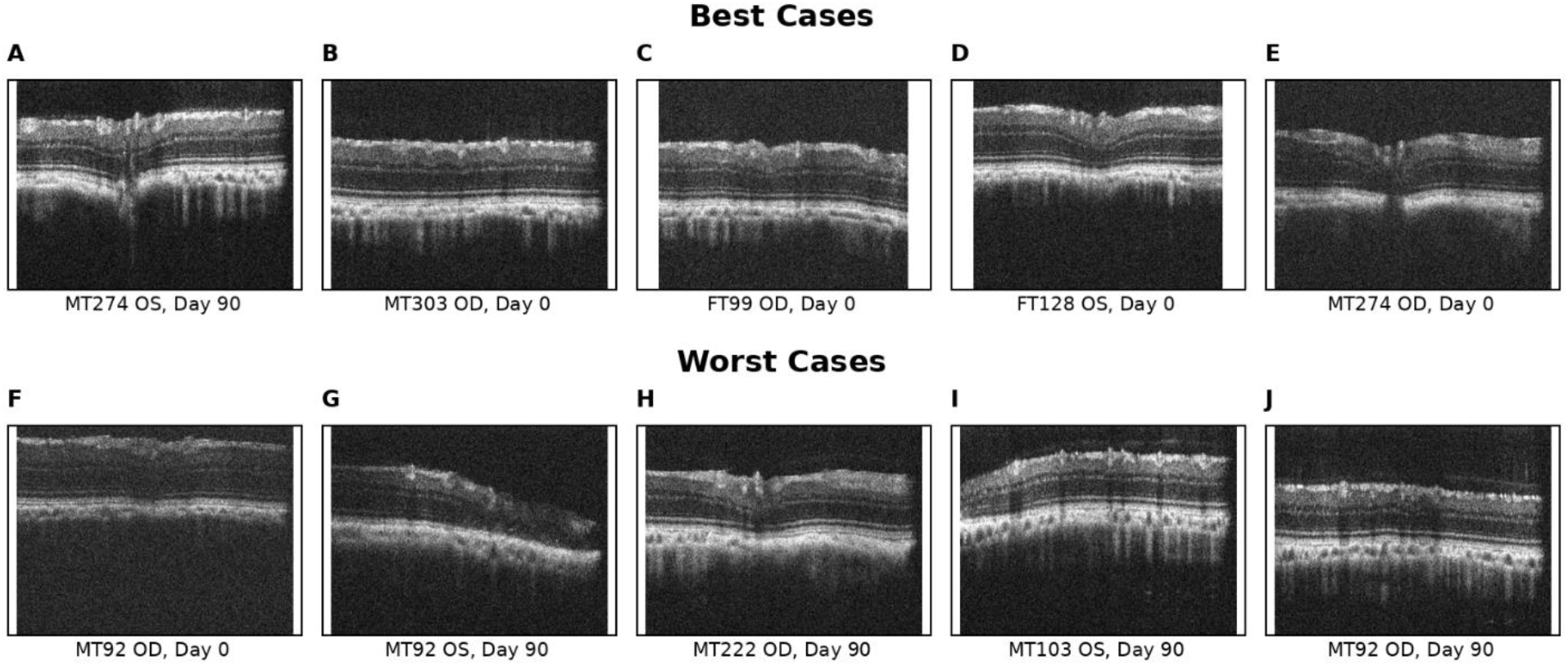
Best- and Worst-Case Examples from Control Test Set. Representative raw OCT B-scans from the control evaluation set. The top row shows five lowest-error cases and the bottom row shows five highest-error cases. The failure cases frequently exhibit reduced contrast, brightness inconsistency, partial cropping, or acquisition artefacts, matching the quality-related failure modes discussed in Methods.

**Figure 2:**
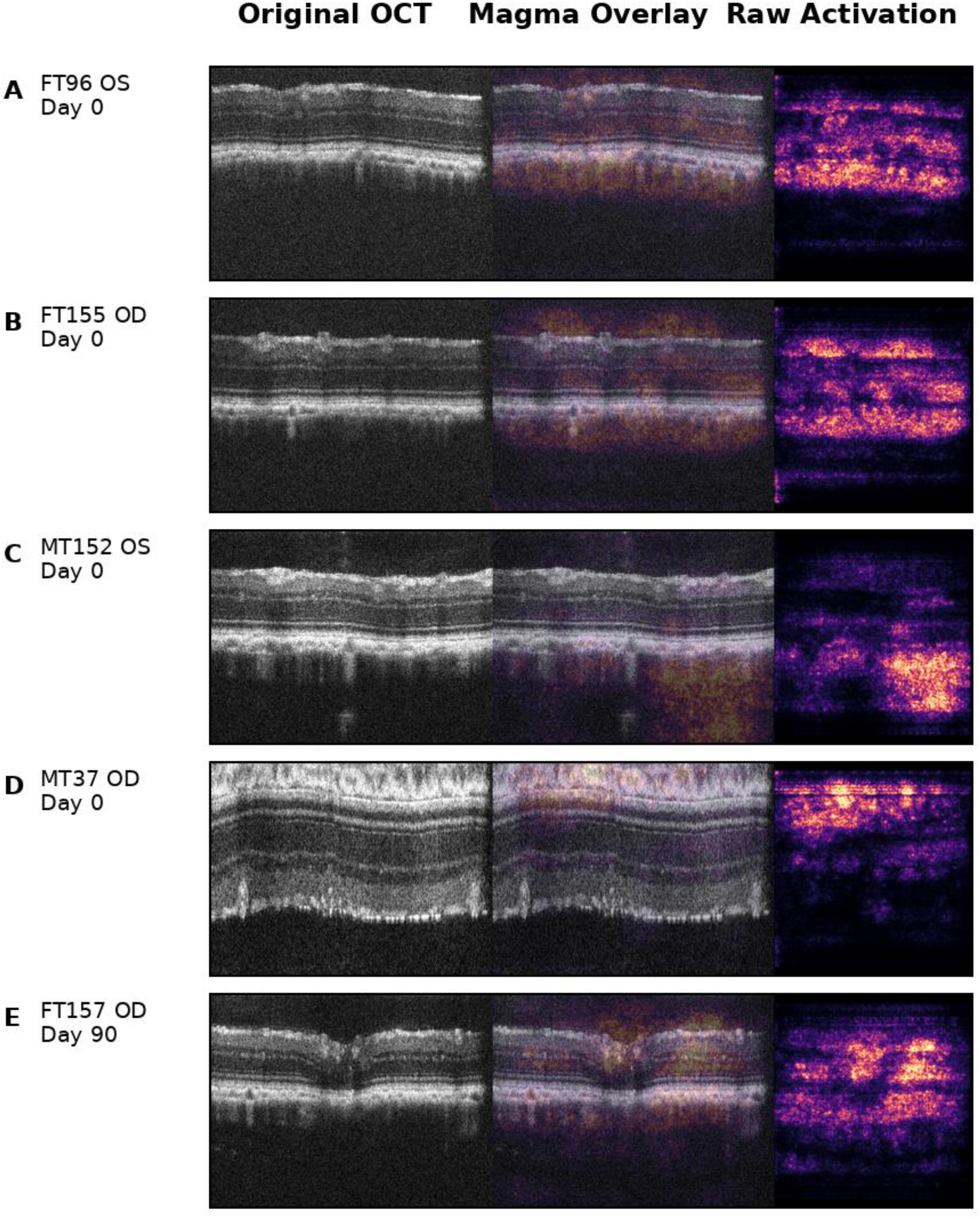
Saliency Map Examples — Anatomically Plausible Attention. Five low-error control examples from the best-performing Xception + GAP model, shown as representative B-scan triplets with the original OCT image, a low-opacity magma saliency overlay, and the raw activation map. High-activation regions localize mainly to the RNFL, optic nerve head-adjacent tissue, and inner retinal layers, supporting anatomically plausible attention patterns while remaining qualitative and hypothesis-generating.

**Figure 3:**
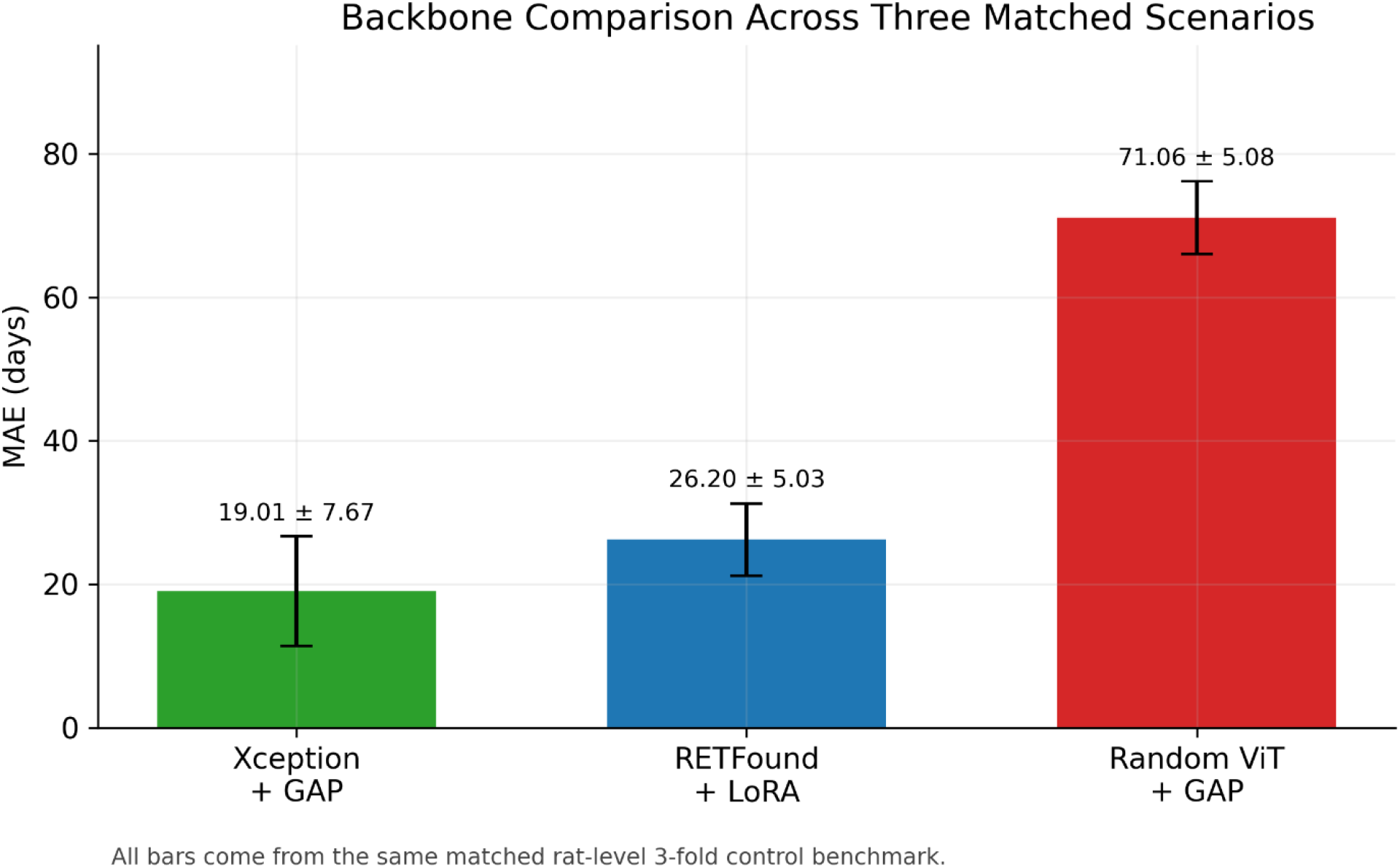
Backbone / Adaptation Comparison Across Four Scenarios. Bar chart comparing control-set MAE across three scenarios: Xception + GAP, RETFound + LoRA and Random ViT + GAP. Error bars indicate standard deviation where available. Xception + GAP, RETFound + LoRA, and Random ViT + GAP come from the matched rat-level 3-fold control benchmark.

**Figure 4:**
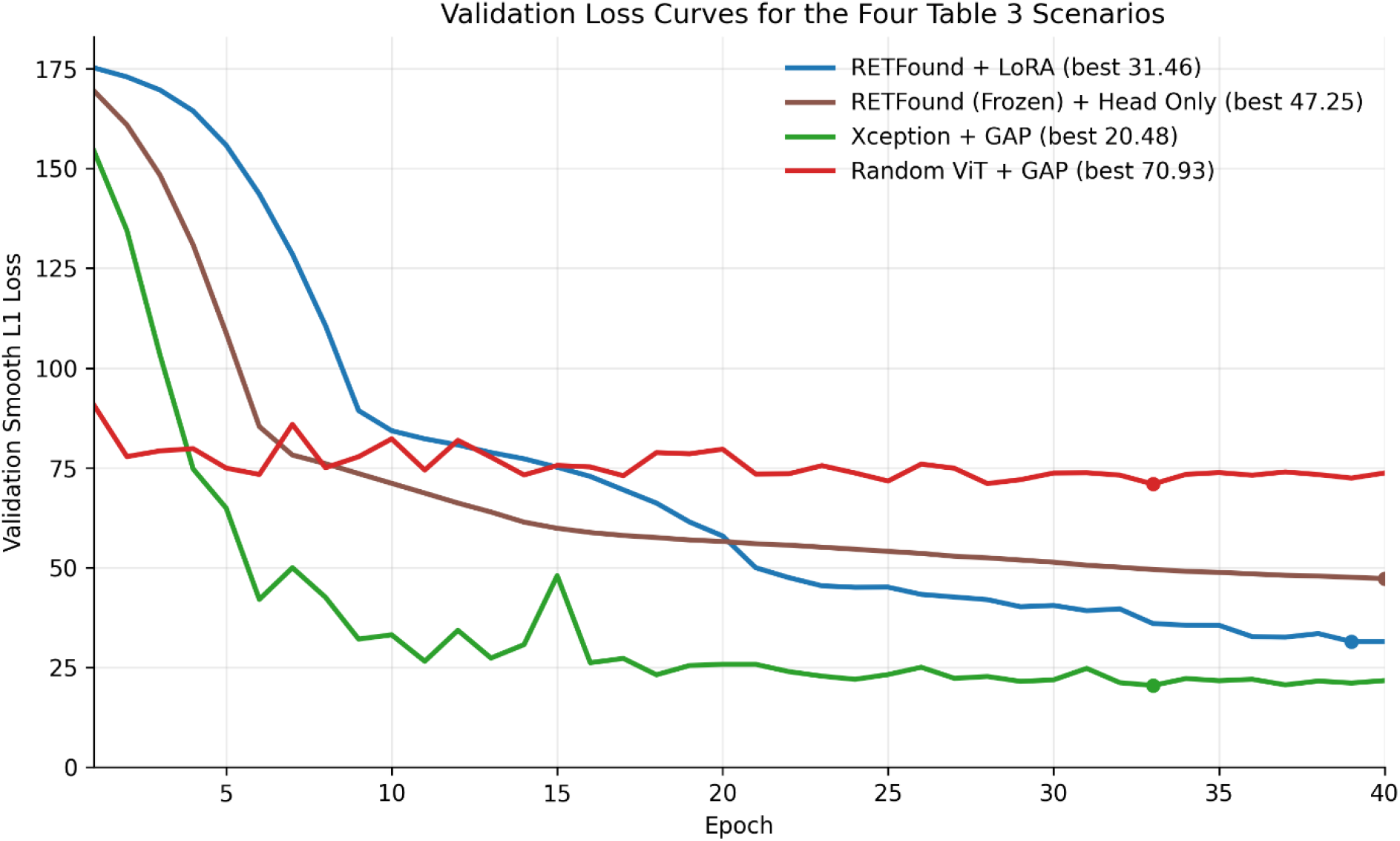
Validation Loss Curves — Comparison of Four Training Scenarios. Validation Smooth L1 loss across 40 epochs for four training scenarios: RETFound + LoRA, RETFound with frozen backbone and trainable head only, Xception + GAP, and a scratch/random ViT negative-control baseline. Xception reached the lowest validation loss, RETFound + LoRA improved steadily but remained above Xception, the frozen-head condition plateaued higher, and the scratch/random ViT remained worst throughout training.

While saliency methods do not prove causality, convergence between reliability metrics and anatomically plausible attention patterns strengthens confidence that the predicted age signals represent structured model responses to retinal features rather than noise.

### 4.1 Principal Findings

This study presents a reproducible benchmark for adapting a human retinal foundation model to rodent OCT age estimation in a data-scarce space-biology setting. Using OSD-679 control animals, RETFound adapted with LoRA achieved MAE = 26.20 ± 5.03 days and R^2^ = 0.744 ± 0.049 under matched 3-fold cross-validation while maintaining inter-eye MAD = 14.22 days. These results indicate that human-pretrained retinal representations remain usable after cross-species transfer, even when the downstream dataset is limited.

The benchmark comparison also yielded an informative finding: a simpler ImageNet-pretrained CNN, Xception, outperformed RETFound on this task (MAE = 19.01 ± 7.67 days; R^2^ = 0.853 ± 0.082). This suggests that foundation-model transfer does not automatically guarantee superior performance in cross-species, small-sample OCT settings. Methodologically, this reinforces the value of matched architecture benchmarking when evaluating foundation models in space-biology imaging tasks.

### 4.2 Foundation Models in Cross-Species Transfer: Benefits and Limits

Human-pretrained retinal features therefore appear useful but not dominant in this setting. Xception likely benefited from a stronger bias toward local texture, layer boundaries, and reflectivity gradients, which are central to OCT interpretation and are known to be effectively captured by convolutional architectures, particularly in small-sample regimes where transformer models may be less data-efficient.[20]

This interpretation fits the anatomy and data regime of the task. Human and Brown Norway rat retinas share a conserved layered retinal organization, providing a plausible structural basis for transferring OCT-derived age-related features across species.[7,8] However, important differences remain: the rat retina lacks a fovea, is rod-dominant, and differs in scale and optic nerve head architecture, as also reflected in prior rodent OCT analysis literature.[24] Rat OCT B-scans are structurally regular, the dataset remains modest, and the target anatomy differs from the human pretraining domain in ways that matter. Under those conditions, stronger convolutional inductive bias may be more valuable than broader model capacity.

The practical implication is straightforward: space-biology AI studies should benchmark architecture families rather than defaulting to the largest available pretrained model. In this setting, empirical comparison was more informative than model pedigree alone. More broadly, the results suggest that cross-species retinal transfer is feasible, but that performance depends strongly on the interaction between anatomy, sample size, and model inductive bias.

### 4.3 Parameter-Efficient Adaptation (LoRA) for Space Medicine

LoRA nevertheless remains valuable as an adaptation strategy. It reduced the number of trainable parameters by approximately 98% relative to full fine-tuning, reached its best validation loss earlier, and preserved a compact update footprint that is easier to store and distribute than a fully tuned backbone. Those are meaningful practical advantages in a small-sample research setting, even though full fine-tuning achieved lower raw control MAE in the adaptation ablation.

The adaptation ablation also clarified a lower bound on model flexibility: frozen head-only training performed worst, indicating that some backbone adaptation is required for effective cross-species transfer. In other words, the task does not support a purely fixed feature extractor, but it also does not guarantee that the most parameter-efficient strategy will be the most accurate.

### 4.4 Reliability and Explainability

Inter-eye agreement supports the internal stability of the models. In the primary RETFound evaluation, inter-eye MAD was 14.22 days (8.56% of mean age), while the matched Xception comparison yielded 16.71 days (10.07%). These values suggest that predictions were driven by reasonably stable anatomical signal rather than by purely idiosyncratic scan noise, although reliability should still be interpreted alongside accuracy rather than in isolation.

Saliency maps further corroborated non-random behavior. Because spaceflight-associated neuro-ocular syndrome (SANS) hypotheses frequently emphasize optic nerve head, RNFL, and pressure- or fluid-sensitive remodeling, the concentration of model attention in these regions is more consistent with biologically meaningful retinal structure than with arbitrary background texture.[10-12] Nevertheless, the saliency maps remain hypothesis-generating only and require direct multimodal validation against physiology, histology, or omics endpoints.

### 4.5 Implications for Space Health AI

This work also leaves a practical roadmap. Future extensions should test the pipeline on additional rodent space-analog conditions, quantify associations with direct ocular endpoints such as IOP or layer thickness, and evaluate whether intermediate domain adaptation or multimodal supervision can narrow the gap between pretrained transformers and strong CNN baselines.

### 4.6 Limitations

Several limitations warrant consideration. Cross-species anatomy differences constrain transferability, cohort composition was imbalanced, and sex was partially confounded with cohort and age range. Cohort 4 was excluded because of limited sample size and design imbalance. The main analyses were intentionally restricted to Day 0 and Day 90 to maximize stability, which limits inference about intermediate longitudinal dynamics. Mechanistic interpretation remains modest because the study does not directly link predicted age to physiological or histological endpoints. Although an HLS external/OOD set was reserved during development, those results were not a focus of the present control-benchmark paper and are therefore not emphasized here. External validation is also incomplete: the results are specific to this SD-OCT setting in OSD-679 and should be tested in additional rodent datasets and, where protocol alignment permits, in human OCT cohorts.

A practical next validation step is two-stage external testing: first across additional rodent space-analog conditions, including recovery intervals and hypercapnia, and then across human OCT datasets where acquisition and annotation protocols are sufficiently aligned to support meaningful comparison.

### 4.7 Conclusion

In summary, we present a cross-species benchmark for adapting a human retinal foundation model to rodent OCT age estimation in a data-scarce space-biology setting. RETFound+LoRA provided useful transfer performance, but a strong pretrained CNN baseline achieved better accuracy under the primary matched benchmark. These findings support the use of empirical architecture benchmarking rather than default assumptions about foundation-model superiority. More broadly, this preprint provides a reproducible baseline for future retinal biomarker development and cross-species transfer studies in space-relevant datasets. As a preprint, we hope this work serves as an openly reusable benchmark that can be extended across additional rodent cohorts, space-analog paradigms, and aligned human OCT datasets.

Table 2 cohort labels reflect the Day 0/90 control subset composition (Cohort 1 = 41 rats, Cohort 2 = 50 rats, and Cohort 3 = 32 rats; Supplementary Table S1), while the MAEs are pooled across out-of-fold eye-day predictions. Cohort 3 also entered the study at an older baseline age (270 days), so its Day 0 and Day 90 rows correspond to 270- and 360-day evaluations rather than the 90- and 180-day regime represented by Cohorts 1 and 2; this likely contributes to the distinct error pattern and should not be interpreted as a simple isolated cohort effect.

Table 3 is restricted to the prespecified main-text backbone comparison between RETFound and Xception on matched 3-fold control splits. The full three-backbone comparison including the scratch/random transformer is reported in Supplementary Table S4, whereas the RETFound adaptation ablation (LoRA vs. full fine-tuning vs. frozen head-only) is reported separately in Supplementary Table S5.

Here, ‘Control’ refers to the non-HLS control arm of OSD-679 rather than a separate nested subgroup. The RETFound and Xception rows summarize out-of-fold control evaluations under their respective matched 3-fold control protocols. Inter-eye MAD is reported as a reliability descriptor and should be interpreted alongside age-prediction accuracy, not as a standalone measure of overall model quality.

## Data and Code Availability

All code, configuration files, split definitions, trained model checkpoints, and the polished OSD-679 age-prediction reproducibility bundle are available at: https://github.com/Alirezahayatimedtech/LoRaRetAgePred

OSD-679 data must be requested via NASA GeneLab: https://genelab-data.ndc.nasa.gov/genelab/accession/OSD-679

## Acknowledgments

We thank NASA GeneLab and the OSD-679 investigators for generating and releasing the dataset.

## Ethics Statement

This study is a secondary analysis of publicly available OCT images from NASA GeneLab OSD-679. All animal procedures were performed under the approvals and oversight described in the original OSD-679 study documentation.

